# CRISPRi-Linked Multi-Module Negative Feedback Loops to Address Winner-Take-All Resource Competition

**DOI:** 10.1101/2025.05.15.654351

**Authors:** Sadikshya Rijal, Kylie Standage-Beier, Rong Zhang, Austin Stone, Abdelrahman Youssef, Xiao Wang, Xiao-Jun Tian

## Abstract

Cellular resource limitations create unintended interactions among synthetic gene circuit modules, compromising circuit modularity. This challenge is particularly pronounced in circuits with positive feedback, where uneven resource allocation can lead to Winner-Takes-All (WTA) behavior, favoring one module at the expense of others. In this study, we experimentally implemented a Negatively Competitive Regulatory (NCR) controller using CRISPR interference (CRISPRi) and evaluated its effectiveness in mitigating WTA behavior in two gene circuits: dual self-activation and cascading bistable switch. We chromosomally integrated a tunable dCas9 gene and designed module-specific gRNAs, with each module encoding its own gRNA to self-repress via competition for limited dCas9. This configuration introduces strong negative feedback to the more active module while reallocating resources to the less active one, promoting balanced module activation. Compared to the control group lacking dCas9-mediated repression, the NCR controller significantly increased module coactivation and suppressed WTA behavior. Our quantitative results demonstrate that NCR provides an effective strategy for regulating resource competition and improving the modularity of synthetic gene circuits.

## INTRODUCTION

The behavior of heterologous gene circuits in host cells remains difficult to predict, largely due to complex circuit-host interactions^1–3^. In particular, when multiple circuit modules coexist, competition for limited cellular resources can lead to unintended and often detrimental outcomes^4–9^. Synthetic constructs not only burden host machinery but also compete with one another for transcriptional and translational resources, reducing each other’s expression by up to 60% and compromising modularity^4,10^. Notably, we previously showed that resource competition is especially severe in circuits driven by positive feedback, where it induces a ‘Winner-Takes-All’ (WTA) effect ^8^. In such systems, resource allocation becomes highly imbalanced, one module dominates while the other is silenced, making it difficult to achieve stable coactivation between co-expressed bistable switches.

Multiple strategies have been developed to regulate circuit-host interaction, including orthogonal resource systems, division of labor via microbial consortia, phase separation and feedback and feedforward control mechanisms^11–15,8,16–25^. For instance, a multi-strain approach has been used to overcome WTA resource competition^8^, though its long-term stability may be compromised by differences in strain growth rates. While various negative feedback and feedforward strategies have effectively stabilized expression in single-module systems^17–25^, extending these approaches to multi-module circuits remains nontrivial. We previously proposed several multimodule control strategies based on negatively competitive regulation (NCR) ^26–29^ and theoretically demonstrated their effectiveness in mitigating WTA resource competition. In these strategies, a single controller resource is deliberately limited and competitively allocated to mediate feedback or feedforward inhibition of each module. As a result, a highly active module is penalized more strongly by the NCR controller while simultaneously depleting the shared controller resource, thereby relieving inhibition on less active modules for its activation.

In this study, we experimentally implemented a CRISPR-driven multimodule negative competitive regulation (NCR) strategy in E. coli, leveraging catalytically deactivated Cas9 (dCas9) as a synthetic, limited resource competed over by module-specific single guide RNAs (sgRNAs). By applying negative feedback through dCas9-mediated transcriptional repression, each module can downregulate itself based on its sgRNA expression. We applied this strategy to enhance coactivation between two bistable switches in both DSA and CBC circuit architectures. Our results show that NCR effectively mitigates WTA behavior and promotes robust coactivation, regardless of WTA direction or circuit topology. This improvement was validated by comparison to a control lacking the NCR controller.

## RESULTS

### Design of CRISPR-Linked Multi-Module Negative Feedback Loops

Figure 1a illustrates the overall design of the CRISPR-linked multi-module negative feedback regulation. The synthetic gene circuit consists of two modules, M1 and M2, encoded on a multi-copy plasmid, while dCas9 is expressed from the bacterial chromosome. To make the CRISPRi system tunable, dCas9 expression is placed under the control of the pTet promoter. The TetR repressor is constitutively expressed (e.g., from the J23117 promoter) on the same plasmid as M1 and M2. This configuration allows the level of dCas9 to be modulated by the addition of anhydrotetracycline (aTc), which inhibits TetR and thereby induces dCas9 expression, enabling controlled negative feedback regulation.

**Figure 1.**
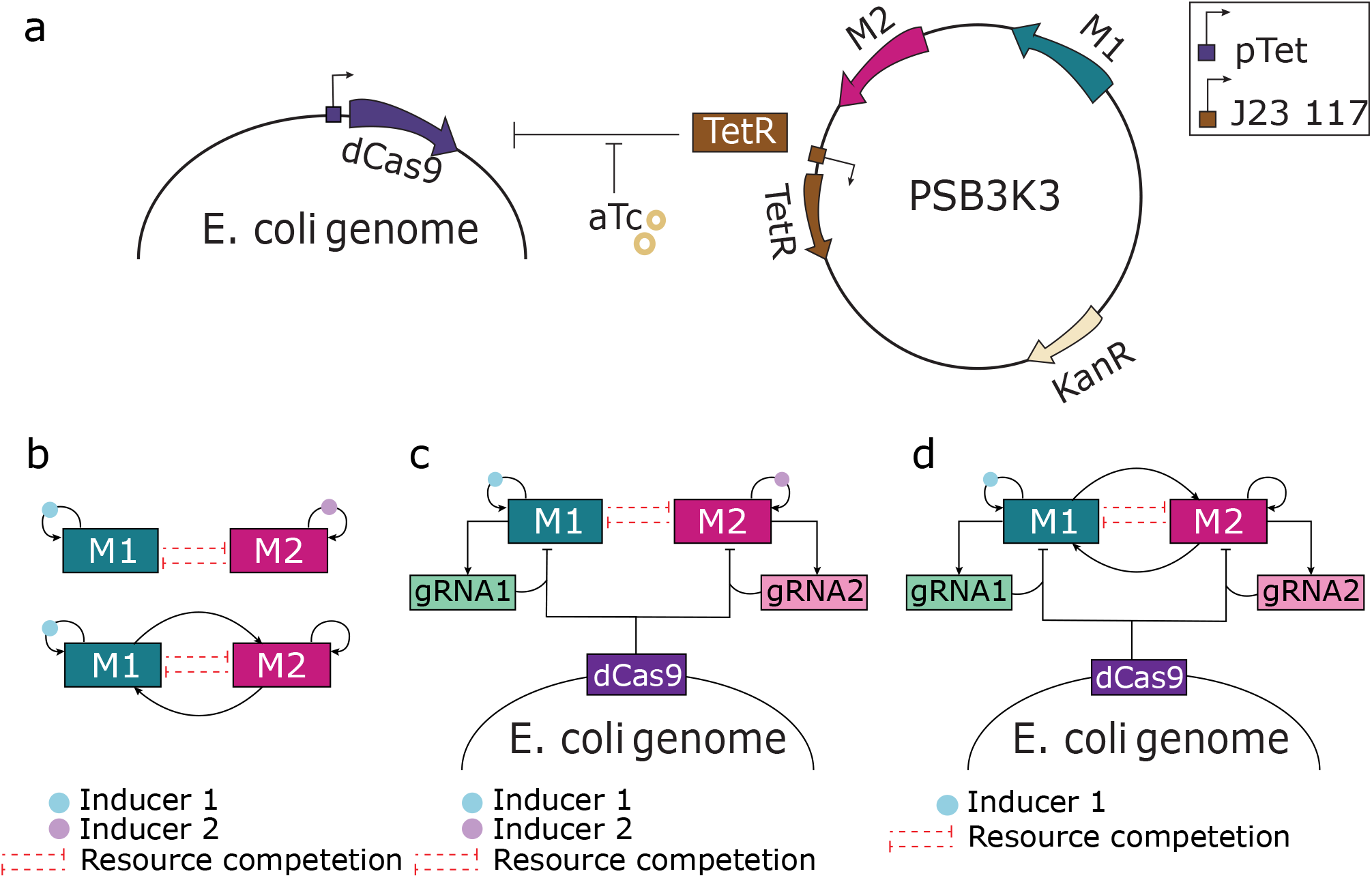
Schematic of the CRISPR-linked multi-module negative feedback design for resource competition mitigation. (**a**) The synthetic gene circuit comprises two modules (M1 and M2) encoded on a multi-copy plasmid, while dCas9 is chromosomally integrated and placed under the aTc-inducible pTet promoter. Constitutive expression of TetR from the same plasmid represses dCas9 expression in the absence of aTc. (**b**) Schematic of the Dual Self-Activation (DSA) circuit (top) and the Cascading Bistable Switch (CBS) circuit (bottom), both exhibiting Winner-Takes-All (WTA) behavior. Each circuit comprises two self-activating modules, M1 and M2, in which the transcription factor of each module promotes its own expression. Red dotted lines between M1 and M2 represent unintended mutual inhibition caused by competition for shared cellular resources. (**c-d**) Implementation of CRISPR-based NCR controller in the DSA circuit (d) and CBS circuit (c). Each module expresses a unique guide RNA (gRNA) that targets its own promoter, enabling module-specific transcriptional repression through competition for a shared, limited pool of dCas9.

Previously, WTA resource competition behavior was observed in two types of gene circuits: the dual self-activation circuit (DSA) and the cascading bistable switch (CBS) (Fig. 1b). Both circuits contain two self-activation modules, M1 and M2, in which each transcription factor activates its own expression. Instead of the expected coactivation, both circuits exhibited counterintuitive WTA behavior, driven by resource competition between the modules. Therefore, implementing the CRISPRi-based NCR controller in DSA and CBS allows us to assess its effectiveness in mitigating WTA behavior in positive feedback circuits.

To implement negative feedback regulation, one guide RNA (gRNA) is designed per module to target its respective promoter (Fig. S1). Each gRNA forms a complex with dCas9 to repress transcription of its own module, thereby closing the negative feedback loop (Fig. 1c, 1d). In this design, the two gRNAs compete for a fixed pool of dCas9, introducing a layer of negatively competitive regulation. This competition counterbalances the intrinsic resource competition between M1 and M2. A module with higher activity produces more gRNA, sequestering more dCas9 and thereby increasing its self-inhibition while relieving inhibition on the other module. This dynamic design benefits the lower-activity module by both freeing up cellular resources and reducing the dCas9-mediated inhibition, promoting more balanced gene expression and resource allocation between the two modules.

### Evaluation of CRISPR-mediated regulation of single modules

To assess the inhibitory efficacy of the chromosome-integrated dCas9 system, we first constructed a dual self-activation (DSA) circuit, DSA-T, comprising an AraC self-activation module (M1) and a LuxR self-activation module (M2) (Fig. 2a). In M1, the transcription factor AraC activates its own expression, along with pBAD-targeting sgRNA (sg-pBAD) and GFP, by binding to the pBAD promoter in the presence of L-arabinose (L-ara). Similarly, in M2, LuxR activates its own expression, the pLux9-targeting sgRNA (sg-pLux9), and RFP via the pLux9 promoter in response to the quorum-sensing signal 3-oxo-C6-HSL (C6). A corresponding control circuit, DSA-C, was also constructed using non-targeting sgRNAs for both modules.

**Figure 2.**
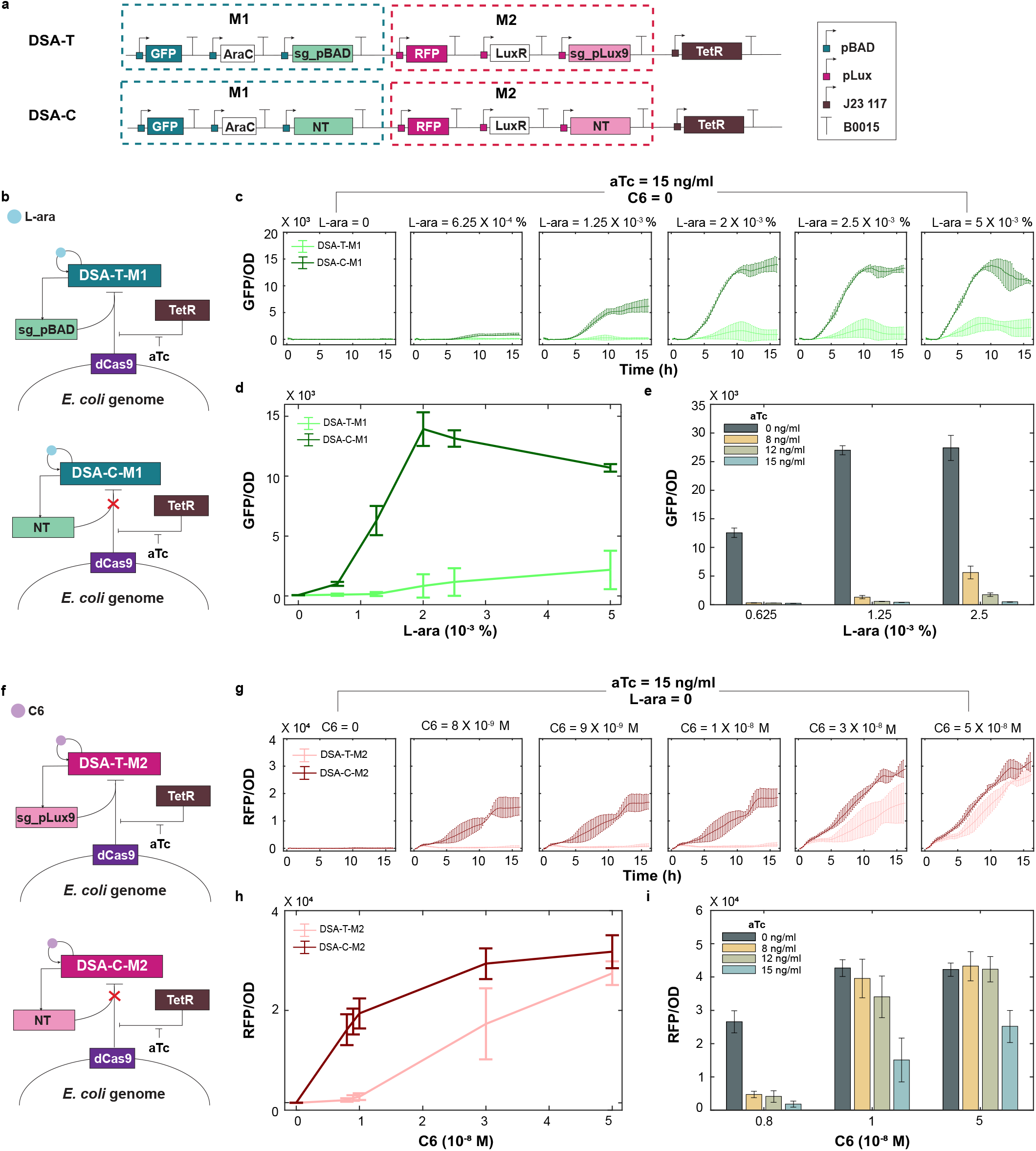
Evaluation of CRISPR Transcriptional Repression in Single Modules. (**a**) Design of the DSA circuit with embedded NCR controller (DSA-T) and its control (DSA-C). Both circuits consist of an AraC self-activation module (M1) and a LuxR self-activation module (M2), induced by L-ara and C6, respectively. Each module co-expresses a promoter-targeting sgRNA (sg-pBAD for M1 and sg-pLux9 for M2) to enable dCas9-mediated negative feedback. The control circuit (DSA-C) expresses non-targeting sgRNAs, disabling negative feedback regulation. (**b**) Network diagrams of CRISPR-mediated transcriptional regulation in module M1 for the test circuit (DSA-T-M1, top) and control (DSA-C-M1, bottom). (**c**) Time-course analysis of GFP/OD in DSA-T-M1 and DSA-C-M1 across increasing concentrations of L-ara, with constant induction of dCas9 by 15 ng/mL aTc. Module M2 remains inactive under 0 nM C6. (**d**) Steady-state dose–response curves of GFP/OD for DSA-T-M1 and DSA-C-M1 as a function of L-ara concentration. (**e**) Steady-state GFP/OD levels of DSA-T-M1 under varying levels of dCas9 interference, controlled by different doses of aTc. (**f**) Network diagrams of CRISPR-mediated transcriptional regulation in module M2 for the test circuit (DSA-T-M2, top) and control (DSA-C-M2, bottom). (**g**) Time-course analysis of RFP/OD in DSA-T-M2 and DSA-C-M2 across increasing concentrations of C6, with constant induction of dCas9 by 15 ng/mL aTc. Module M1 remains inactive under 0 nM C6. (**h**) Steady-state dose–response curves of RFP/OD for DSA-T-M2 and DSA-C-M2 as a function of C6 concentration. (**i**) Steady-state RFP/OD levels of DSA-T-M2 under varying levels of dCas9 interference, controlled by different doses of aTc. Data in the curve represents means ± s.d. n=3.

To evaluate the repression efficiency of CRISPRi on gene expression in M1, we performed a dose-response analysis using DSA-T-M1 and DSA-C-M1 circuits (Figure 2b) across a range of L-ara concentrations, with M2 uninduced. Under fixed dCas9 expression (15 ng/mL aTc), we monitored GFP fluorescence over time following induction with varying L-ara concentrations. As shown in Figure 2c, the DSA-T circuit exhibited slower and weaker activation compared to the control DSA-C circuit. The steady-state dose-response curve revealed significant GFP repression in the DSA-T circuit compared to the control DSA-C circuit (Figure 2d). To assess tunability, we varied aTc concentrations and observed consistent fluorescence downregulation, with reduced repression at 8 ng/mL and strong repression at 15 ng/mL, demonstrating aTc-dependent control of CRISPRi strength (Figure 2e).

A similar analysis was performed to evaluate CRISPRi-mediated repression of M2 in the DSA-T-M2 and DSA-C-M2 circuits across a range of C6 concentrations, with L-ara absent (Figure 2f). With aTc fixed at 15 ng/mL, time-series data showed weaker and delayed RFP expression in the test circuit (DSA-T-M2) compared to the control (DSA-C-M2) at moderate C6 levels (Figure 2g). However, at higher C6 doses, repression was less effective, suggesting asymmetric CRISPRi efficiency between M1 and M2. This trend was confirmed by the steady-state dose-response curve (Figure 2h) and further supported by fluorescence data collected at varying aTc levels (Figure 2i). The reduced repression at high C6 concentrations is likely due to competition between CRISPRi (via sg-pLux9) and the LuxR–C6 complex for promoter binding, as the sgRNA target site in pLux9 overlaps with the Lux box.

### Mitigation of WTA resource competition in Dual Self-Activation (DSA) circuit

We implemented the CRISPR-based NCR controller in the DSA circuit to evaluate its effectiveness in multimodule regulation and mitigation of WTA behavior. First, we used Flow cytometry to analyze cell fate transitions in DSA-T and DSA-C circuits by varying the M2 inducer (C6) while keeping the M1 inducer (L-ara) fixed at 1.25 × 10^−3^%. The aTc concentration was maintained at 15 ng/mL to ensure effective dCas9-mediated repression. In the control circuit (DSA-C), pronounced WTA behavior was observed under imbalanced induction. As shown in Fig. 3a (top), cells shifted from an M1-only state (GFP-high/RFP-low) at low C6, to a transient coactivation state (GFP-high/RFP-high) at intermediate C6, and then to an M2-only state (GFP-low/RFP-high) at high C6, consistent with previous results^8^. In contrast, in the system with DSA-T circuit, M1 was initially suppressed due to dCas9 inhibition, but activation of M2 by C6 also promoted M1 activation (Fig. 3a, bottom). This occurs due to the designed cross-activation, where activation of one module sequesters more dCas9, thereby reducing repression on the other. Notably, DSA-T system showed a substantial increase in coactivation at higher C6 concentrations (Fig. 3a, bottom), where WTA dominated in the control system. Quantitative analysis of flow cytometry data showed that in the DSA-C circuit, the coactivation fraction initially increased with C6 induction but later declined as more cells transitioned into the M2-only WTA state. In contrast, the DSA-T circuit exhibited a steady increase in coactivation with rising C6 levels (Fig. 3b-c). This strongly suggests that our NCR controller prevents M2 from monopolizing resources by repressing its transcription and redistributing resources to support M1 expression.

**Figure 3.**
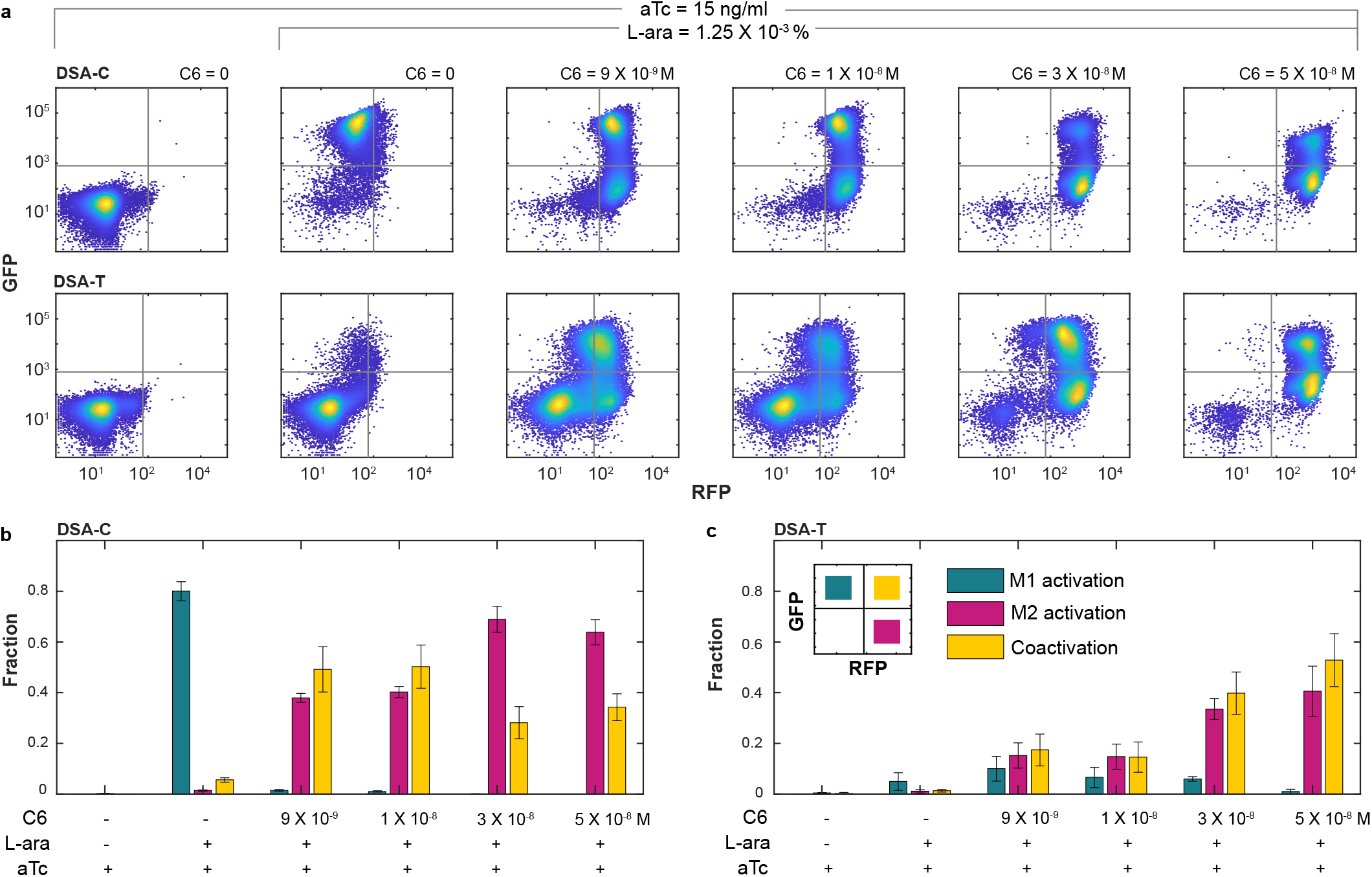
Mitigation of WTA resource competition in DSA circuits. (**a**) Flow cytometry data illustrate cell state transitions in DSA-C (top) and DSA-T (bottom) with increasing C6 concentrations, a fixed L-ara dose (1.25 × 10^−3^ %), and constant aTc (15 ng/mL). For each sample, 10,000 events were recorded. The data shown represent one of three independent biological replicates. (**b-c**) The fraction of cells in M1 activation (green), M2 activation (pink), and coactivation (yellow) states in DSA-C (c) and DSA-T (d) across increasing C6 concentrations. Data are presented as mean ± s.d., n = 3.

To further validate the WTA mitigation efficacy of the CRISPRi-based NCR controller, we examined cell fate transitions by increasing L-ara while maintaining a constant C6 concentration in both DSA-C and DSA-T circuits. The aTc concentration was fixed at 6 ng/mL, as M1 is particularly sensitive to dCas9-mediated repression. As shown in Supplementary Fig. S2, the DSA-C circuit displayed a typical WTA pattern, with high L-ara levels driving cells into an M1-only WTA state following a brief coactivation phase at intermediate L-ara doses. In contrast, the DSA-T circuit maintained a stable coactivation population across the L-ara gradient. Together, these findings demonstrate that the dCas9-mediated multimodule NCR strategy robustly suppresses WTA behavior in both directions of cell fate transitions in the DSA circuit.

### Mitigation of WTA resource competition in the synthetic cascading bistable switches (Syn-CBS) circuit

To assess the generalizability of the CRISPR-based NCR controller, we implemented it in a cascading bistable switch (CBS) circuit and compared its performance to a control system. Similar to the DSA circuit, the CBS circuit contains AraC self-activation module (M1) and LuxR self-activation module (M2). However, in CBS, the modules also activate each other (Fig. 4a). Module M1 includes an additional gene, LuxI, which drives the synthesis of the quorum-sensing signal C6 to activate M2, while Module M2 expresses an AraC gene designed to promote M1. To incorporate the NCR controller, we introduced two gRNAs, sg-pBAD and sg-pLux9, targeting M1 and M2, respectively. A corresponding control circuit (CBS-C) was constructed using non-targeting gRNAs, thereby eliminating CRISPR-mediated repression (Fig. 4b)

**Figure 4.**
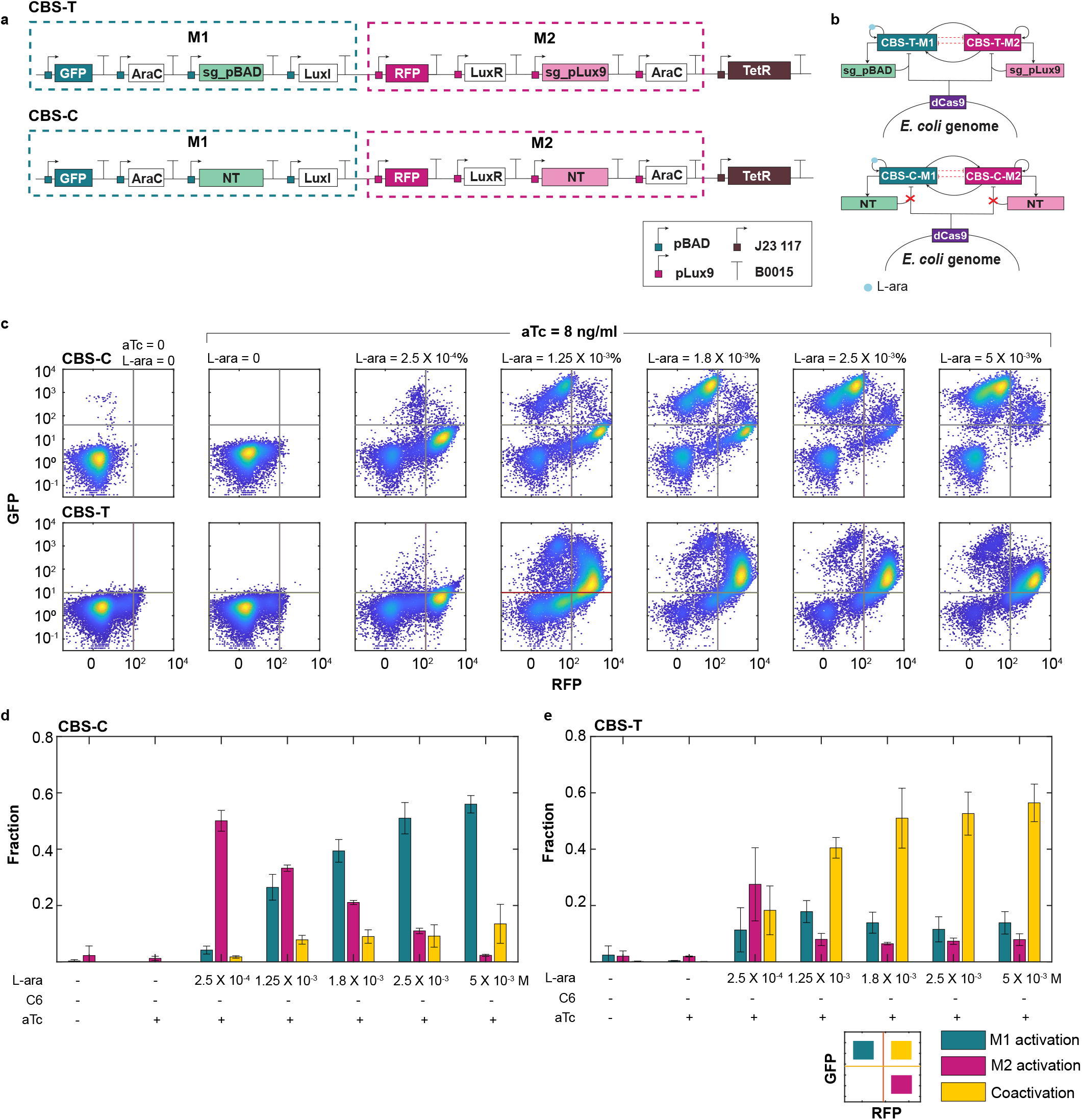
CRISPR-driven WTA mitigation in CBS circuits. (**a**) Design of the cascading bistable switch (CBS) circuit with embedded NCR controller (CBS-T) and its corresponding control (CBS-C). Both CBS circuits consist of two self-activation modules, AraC (M1) and LuxR (M2), induced by L-ara and C6, respectively. Module M1 additionally expresses luxI, which synthesizes C6 to activate M2, while Module M2 expresses araC to promote M1 activation. Each module co-expresses a promoter-targeting sgRNA (sg-pBAD for M1 and sg-pLux9 for M2) to enable dCas9-mediated negative feedback in CBS-T. The control circuit (CBS-C) expresses non-targeting sgRNAs, thereby lacking negative feedback regulation. (b) Network diagrams of the circuit CBS-C (top) and its control CBS-T (bottom). (**c**) Flow cytometry data illustrate cell state transitions in CBS-C (top) and CBS-T (bottom) with increasing L-ara concentrations and constant aTc (8 ng/mL). For each sample, 10,000 events were recorded. The data shown represent one of three independent biological replicates. (**d-e**) The fraction of cells in M1 activation (green), M2 activation (pink), and coactivation (yellow) states in CBS-C (d) and CBS-T (e) across increasing Lara concentrations. Data are presented as mean ± s.d., n = 3.

Flow cytometry data across varying L-ara concentrations were analyzed to compare cell fate transitions between CBS-T and CBS-C, with aTc held constant at 8 ng/mL. As shown in Fig. 4c (top), the control system CBS-C exhibited a pronounced shift from an M2-only activation state to an M1-only activation state as L-ara levels increased, with minimal coactivation observed. This strong WTA behavior aligns with our previous findings^8^. In contrast, CBS-T showed a markedly enhanced and stable coactivation population of M1 and M2 (Fig. 4c, bottom), indicating effective mitigation of WTA behavior by the NCR controller.

Cell fate fractions estimated from flow cytometry data revealed a clear WTA pattern in CBS-C: as L-ara concentration increased, the M2-activated population declined, the M1-activated population rose, and coactivation remained minimal (Fig. 4d). In contrast, CBS-T exhibited a steadily increasing coactivation fraction with rising L-ara levels, eventually reaching saturation (Fig. 4e). Similar trends were observed at alternative aTc concentrations, suggesting that modest changes in dCas9 induction do not compromise NCR function (Supplementary Fig. S3). Overall, these results demonstrate that the NCR controller prevents abrupt cell fate transitions from M2 to M1-dominant states, mitigates WTA resource competition, and promotes balanced expression between M1 and M2.

## DISCUSSION

The loss of modularity in heterologous gene circuits, driven by unintended host-circuit and circuit-circuit interactions, is a pervasive challenge in synthetic biology^1^. To address this, a variety of control strategies have been developed to construct resource-aware systems^1,2^. In this study, we demonstrated that resource competition between two self-activating bistable switches can be effectively mitigated using negatively competitive regulation (NCR) mediated by CRISPR-based transcriptional repression. We validated the generalizability of this approach through experiments on both the DSA and CBS constructs. The NCR controller enables efficient multi-module regulation with minimal cellular burden. This is achieved by maintaining low expression of chromosomally integrated dCas9 and leveraging sgRNAs, which do not rely on ribosomes for expression. Together, these features minimize consumption of host resources and are critical for the optimal performance of the NCR controller.

CRISPR-mediated resource regulation has been widely used to counteract various facets of resource competition in synthetic biology^17,32,33^. Ceroni et. al. introduced CRISPR-driven negative feedback to relieve the metabolic burden of heterologous gene circuits, thereby enhancing host cell growth^17^. Competition among gRNAs for dCas9 can significantly compromise the independent regulation of multiple targets in CRISPR-based systems^30,31,34^. This challenge can be addressed by implementing a dCas9 concentration controller that uses negative feedback on dCas9^32^. A recent theoretical study using a resource-aware model of CRISPR regulation suggests that combining CRISPR interference and activation can help mitigate crosstalk arising from resource competition^33^. Here, we leveraged competition among sgRNAs for limited dCas9 to develop a regulatory strategy that promotes coactivation between two self-activating gene modules. Together, these findings underscore the versatility of CRISPR-based controllers in navigating cellular and/or controller resource limitations to achieve more robust and predictable gene circuit behavior. However, this approach also introduces a trade-off: the negative feedback inherent in the system can attenuate circuit activation and raise the threshold required to trigger state transitions, potentially making the system less responsive to inputs. Future work could focus on optimizing the feedback strength or incorporating dynamic control mechanism to balance responsiveness with robustness in multi-module circuit designs. For example, integrating feedback and feedforward control through recombinase-based promoter flipping has been shown to outperform single negative feedback controllers in resource decoupling^25^.

Efforts toward predictive gene circuit design continue to expand the capabilities of synthetic biology by enabling the construction of increasingly complex systems. In addition to practical applications, these control strategies provide powerful tools for probing the underlying principles of biological processes regulated by intricate gene networks. For example, the epithelial-to-mesenchymal transition (EMT) has been shown to follow a cascading bistable switch (CBS) mechanism^35,36^. In previous work, we employed a division-of-labor approach using two *E. coli* strains to implement synthetic sequential cell fate transitions^8^. However, it is plausible that natural systems have evolved additional negative feedback loops to ensure robust sequential transitions within a single cell. In this study, we demonstrate that the CRISPR-based NCR enables successful coactivation of the synthetic CBS in a single strain. This approach offers multiple advantages: it can provide mechanistic insight into the role of negative feedback in EMT-like transitions, and it promotes minimal heterogeneity and greater long-term stability. The latter is particularly important, as growth rate differences in multi-strain systems often lead to outcompetition and eventual loss of slower-growing, high-burden strains.

## MATERIALS AND METHODS

### Strains, media, and chemicals

We used E. coli strain DH10B (Invitrogen, USA) for cloning, and all the fluorescence measurements were performed in E. coli strain K-12 MG1655ΔlacIΔaraCBAD with catalytically deactivated Cas9 integrated in the chromosome. LB broth (Luria-Bertani broth, Sigma-Aldrich) and LB plates supplemented with 50 μg/ml kanamycin or 100 μg/ml ampicillin, depending on the plasmid backbone, were used for cell growth. The cell colony harboring the plasmid was inoculated in 5 mL LB media with the appropriate antibiotic and grown in a shaking incubator at 250 revolutions per minute for plasmid extraction. We dissolved L-ara (L-(+)-Arabinose, Sigma-Aldrich), C6 (3oxo-C6-HSL, Sigma-Aldrich), and aTc (Anhydrotetracycline hydrochloride, Abcam) in ddH_2_O to prepare stock concentrations of 25%, 10 mM, and 1 mg/ml, respectively. Induced stocks were stored at -20 °C in aliquots to prevent multiple thawing/freezing cycles. L-ara and C6 were diluted in ddH_2_O to working concentrations monthly and were stored at 4 °C. The aTc stock was replaced monthly to ensure efficiency, and the working aTc solutions were prepared freshly each time. All the working solutions were added to the culture media with a 1000-fold dilution for the fluorescence assays. All oligo DNAs were synthesized by Integrated DNA Technologies, Inc.

### Plasmids construction

Unless mentioned otherwise, standard BioBrick parts from the iGEM registry were used for promoters, GOIs, and terminators (Supplementary Table 1). The circuit constructions (Supplementary Table 2) for BioBrick parts were done using BioBrick assembly sites *EcoRI, XbaI, SpeI*, and *PstI* digestions (Respective enzymes from ThermoFisher), followed by dephosphorylation with rSAP phosphatase (Sigma-Aldrich) when necessary, and ligations using T4 DNA ligase (New England BioLabs). Medium copy number (20-30 copies) pSB3K3 backbone with kanamycin resistance was used for circuit construction. The cloned inserts were verified via colony-PCR using insert-specific primers, followed by Sanger sequencing. Gel extraction, PCR reaction cleanup, and plasmid isolation were performed using the GelElute Gel Extraction Kit, GenElute PCR Cleanup Kit, and GenElute Plasmid Miniprep Kit (all from Sigma-Aldrich), respectively. LuxRG2C, AraC, and pLux9 were designed as described previously ^8^. The CRISPR gRNA scaffold was retrieved from pSG4K5 (Addgene, Cat# 74492) using a forward primer to add *XbaI* and *SapI* sites and the reverse primer to add the BioBrick restriction sites to the scaffold. Four additional bases (TGAC) were added to the upstream of pLux9 to ensure a 20-bp binding of sgRNA to the promoter. Similarly, a reverse primer is used to remove additional bases after the TSS (+1) in the pbad promoter and to add *SapI* (New England BioLabs), *SpeI*, and *PstI* sites to it. The pBad promoter plasmid was digested at the *SpeI* site, and the guide RNA scaffold plasmid at the *XbaI* site. The parts are ligated to create the pBad-NT backbone featuring two *SapI* sites for scarless ligation of sg-pBAD. Likewise, forward and reverse oligonucleotides, as mentioned in Supplementary Table 3, were annealed and cloned to *XbaI* and *SapI* sites of pBAD + NT to construct pLux9 + NT. Restriction digestion and scarless ligation using *SapI* sites were performed to build pLux9 + sg-Plux9. A reverse primer with lva-tag was used to add lva-tag to the RFP sequence. An additional H840A mutation was added to pCas9(D10A)(Addgene, Cat# 74495) as described before^37^. Forward and reverse primers with BioBrick sites are used to retrieve the dCas9 for further cloning and genome integration. All the DNA oligos used for the study are listed in Supplementary Table 3.

### Chromosome integration of dCas9

The NCR cassette was integrated into the chromosome using the Clonetegration method as described before^38^. Plasmid pOSIP-KH (Addgene, Cat #45983) (Gift from Drew Endy & Keith Shearwin) was used to integrate dCas9 expressed under pTet promoter into the HK022 site of *E. coli K12* MG1655 (ΔLacI ΔAraC). The dCas9 was cloned into the *EcoR*I and *PstI* sites of plasmid pOSIP-KH, and the ligation mixture was transformed into the destination strain. LB plates supplemented with kanamycin (50 μg/ml) was used for antibiotic screening of positive colonies. Colony PCR was done using dCas9-specific primers to confirm the integration of dCas9 into the chromosome. The positive colonies were made competent and transformed with pE-FLP (Addgene, Cat #45978) (Gift from Drew Endy & Keith Shearwin) to remove the kanamycin resistance cassette from the bacterial chromosome. The pE-FLP positive colonies were selected using LB plates with ampicillin, grown overnight at 30 °C. Selected colonies were inoculated in kanamycin-supplemented LB plates and the removal of kanamycin resistance was confirmed by the absence of the cell growth. Finally, pE-FLP positive colonies were grown overnight in LB media at 37 °C for the plasmid expulsion. E. coli colonies harboring dCas9 in the genome and no antibiotic resistance were selected by streaking the cells in LB plates, followed by negative selection using LB plates supplemented with ampicillin.

### Circuit induction for fluorescence analyses

To collect the plate reader and flow cytometry fluorescence data of GFP and RFP, we began the experiment by transforming pSB3K3 plasmids containing the desired gene constructs into *E. coli*^dcas9^. The transformed culture was grown on kanamycin-supplemented LB plates overnight at 37 °C. Three colonies each from the test and control plates were inoculated into 300 μl of LB medium with 50 μg/ml kanamycin and were grown for 5 hrs. We prepared a 96-well plate of LB medium containing 200 μl of kanamycin-supplemented LB with appropriate inducer concentrations. 5 μL of the cells were inoculated into each well of the 96-well plate and were incubated for 16 h at 37 °C in shaking conditions for fluorescence assays.

### Plate reader average fluorescence analysis

96-well plates inoculated with cells were incubated on Synergy H1 Hybrid Reader (BioTek) at 37 °C with 807 CPM rotational speed. Optical density (OD) was measured as absorbance at 600 nm. GFP and RFP fluorescence were measured at 485/515 nm and 546/607 nm excitation/emission wavelengths, respectively. OD and fluorescence data were analyzed, and plots were generated with MATLAB (R2024b, MathWorks).

### Flow cytometry

Flow data were recorded using Stratedigm S1000EON Flow Cytometer with excitation/emission filters 480 nm/530 nm for GFP and 480 nm/>670 nm for RFP detection. Samples were collected in three biological replicates, and 10,000 events were recorded for each sample. Data files were analyzed with MATLAB (R2024b, MathWorks). Cells were gated using FSC-A/FSC-H to eliminate the doublets and non-cellular small particles with the plain LB medium without any cells as a negative control.

## Data availability

All data produced or analyzed for this study are included in the article and its Supplementary Information files.

## Acknowledgments

This work was supported by grants from the US National Institutes of Health (R35GM142896 to X.-J.T.).

## Author contributions

X.-J.T. conceived the study. X.-J.T., X.W., S.R., R.Z., A.S., and K.S. designed the study. S.R. and A.Y. performed experiments. X.-J.T., S.R., R.Z., and A.S. analyzed the data. X.-J.T., S.R., and A.S. wrote the manuscript. X.-J.T., S.R., and A.S. edited the manuscript.

