## Supplementary Information for "CRISPRi-Linked Multi-Module Negative Feedback Loops to Address Winner-Take-All Resource Competition"

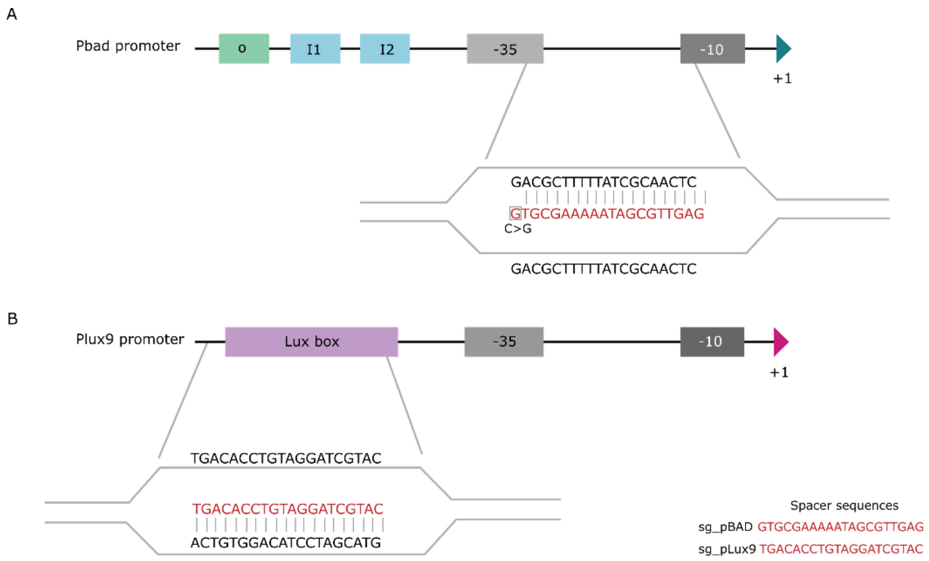


**Figure S1. Design of guide RNAs targeting pBAD and pLux9 promoters.** (**a**) Schematic of the pBAD promoter. Operator sites O, I1, and I2 are critical for AraC-mediated regulation. The -35 and -10 regions are shown as gray boxes, and the transcription start site (TSS) is marked by a green triangle. The sgRNA spacer (red) targets the region between the -35 and -10 boxes and includes a C>G mutation at the 5′ end adjacent to the PAM sequence. (**b**) Schematic of the pLux9 promoter. The -35 and -10 regions are also shown in gray, with the TSS indicated by a pink triangle. The sgRNA spacer (red) is designed to target a region overlapping the LuxR binding site (Lux box).

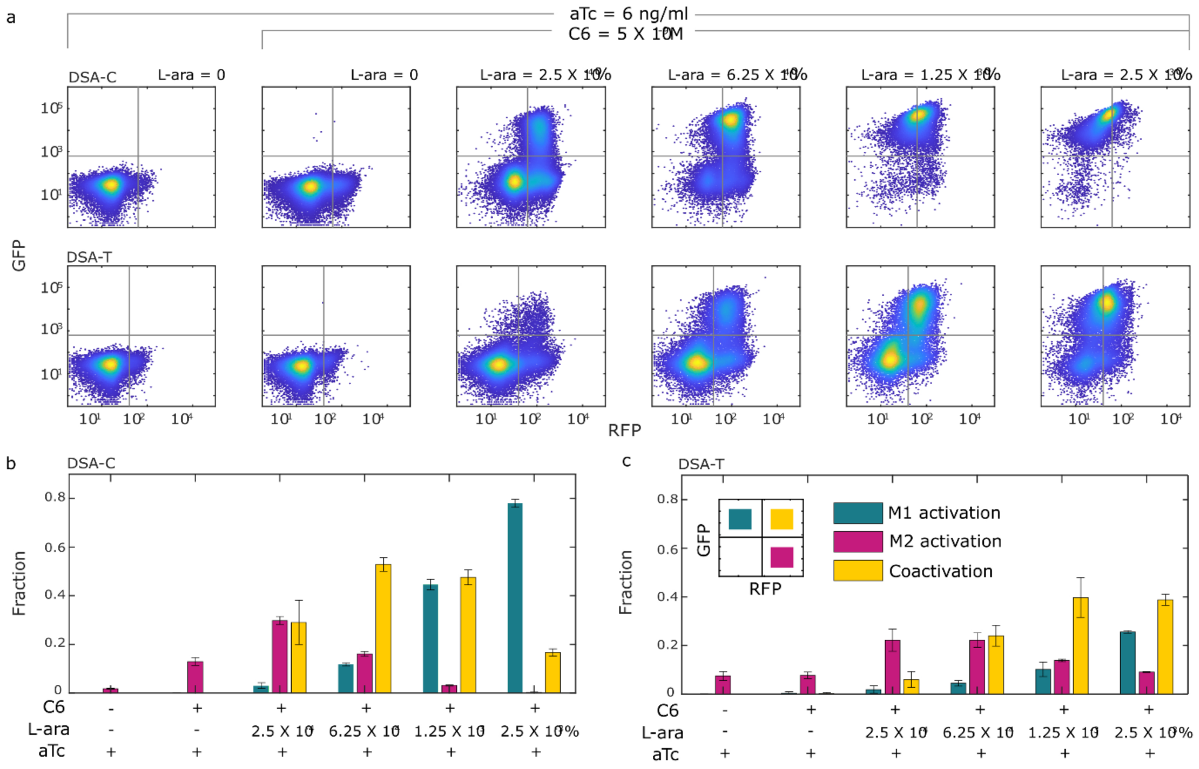


**Figure S2. Mitigation of WTA resource competition in DSA circuits at fixed C6 dose and increasing L-ara**. (a) Flow cytometry data illustrate cell state transitions in DSA-C (top) and DSA-T (bottom) with increasing L-ara concentrations, a fixed C6 (5 X $10^{-9}$ M), and constant aTc (6 ng/mL). For each sample, 10,000 events were recorded. The data shown represent one of three independent biological replicates. (b-c) The fraction of cells in M1 activation (green), M2 activation (pink), and coactivation (yellow) states in DSA-C (c) and DSA-T (d) across increasing L-ara concentrations. Data are presented as mean ± s.d., n = 3.

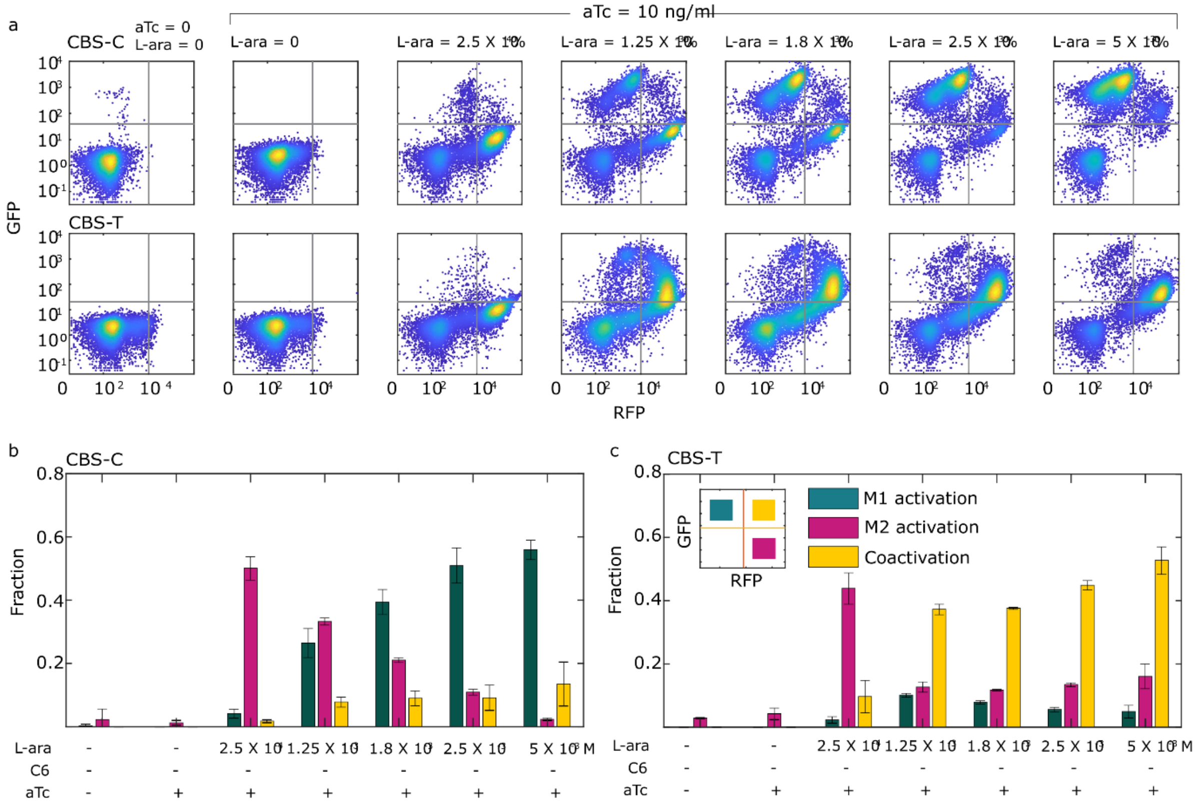


**Figure S3. CRISPR-driven WTA mitigation in CBS circuits at a different level of dCas9.** (a) Flow cytometry data illustrate cell state transitions in CBS-C (top) and CBS-T (bottom) with increasing L-ara concentrations and constant aTc (10 ng/mL). For each sample, 10,000 events were recorded. The data shown represent one of three independent biological replicates. (b-c) The fraction of cells in M1 activation (green), M2 activation (pink), and coactivation (yellow) states in CBS-C (d) and CBS-T (e) across increasing Lara concentrations. Data are presented as mean ± s.d., n = 3.

**Supplemental Table 1. List of BioBrick parts and their sequences used in the study**

| BioBrick codes *(mutations if any)* | Part description | Sequence used in the study (5'→3') |
|---|---|---|
| K206000 *(deletions of bases after the TSS)* | Strong pBad promoter | acattgattatttgcacggcgtcacactttgctatgccatagcaag<br>atagtccataagattagcggatcctacctgacgctttttatcgcaa<br>ctctctactgtttctccataccgtttttttgggctagc |
| R0062 *(A→T at -10 and G→T -11 relative to the TSS)* | A variant of pLux promoter – pLux9 | acctgtaggatcgtacagggttacgcaagaaaatggtttgttata<br>gtcgaataaa |
| P0440 | pTet promoter | tccctatcagtgatagagattgacatccctatcagtgatagagat<br>actgagcac |
| J23117 | J23117 constitutive promoter | ttgacagctagctcagtcctagggattgtgctagc |
| J04031 | Green Fluorescent Protein (GFP) with Lva tag | atgcgtaaaggagaagaacttttcactggagttgtcccaattctt<br>gttgaattagatggtgatgttaatgggcacaaattttctgtcagtg<br>gagagggtgaaggtgatgcaacatacggaaaacttaccctta<br>aatttatttgcactactggaaaactacctgttccatggccaacact<br>tgtcactactttcggttatggtgttcaatgctttgcgagatacccag<br>atcatatgaaacagcatgactttttcaagagtgccatgcccgaa<br>ggttatgtacaggaaagaactatatttttcaaagatgacgggaa<br>ctacaagacacgtgctgaagtcaagtttgaaggtgatacccttg<br>ttaatagaatcgagttaaaaggtattgattttaaagaagatggaa<br>acattcttggacacaaattggaatacaactataactcacacaat<br>gtatacatcatggcagacaaacaaaagaatggaatcaaagtt<br>aacttcaaaattagacacaacattgaagatggaagcgttcaac<br>tagcagaccattatcaacaaaatactccaattggcgatggccct<br>gtccttttaccagacaaccattacctgtccacacaatctgcccttt<br>cgaaagatcccaacgaaaagagagaccacatggtccttcttg<br>agtttgtaacagctgctgggattacacatggcatggatgaactat<br>acaaaaggcctgctgcaaacgacgaaaactacgctttagtag<br>cttaa |
| E1010 | Red Fluorescent Protein (RFP) | atggcttcctccgaagacgttatcaaagagttcatgcgtttcaaa<br>gttcgtatggaaggttccgttaacggtcacgagttcgaaatcga<br>aggtgaaggtgaaggtcgtccgtacgaaggtacccagaccg<br>ctaaactgaaagttaccaaaggtggtccgctgccgttcgcttgg<br>gacatcctgtccccgcagttccagtacggttccaaagcttacgtt<br>aaacacccggctgacatcccggactacctgaaactgtccttcc<br>cggaaggtttcaaatgggaacgtgttatgaacttcgaagacggt<br>ggtgttgttaccgttacccaggactcctccctgcaagacggtga<br>gttcatctacaaagttaaactgcgtggtaccaacttcccgtccga<br>cggtccggttatgcagaaaaaaaccatgggttggaagcttcc<br>accgaacgtatgtacccggaagacggtgctctgaaaggtgaa<br>atcaaaatgcgtctgaaactgaaagacggtggtcactacgac<br>gctgaagttaaaaccacctacatggctaaaaaaccggttcagc<br>tgccgggtgcttacaaaaccgacatcaaactggacatcacctc<br>ccacaacgaagactacaccatcgttgaacagtacgaacgtgc<br>tgaaggtcgtcactccaccggtgcttaa |

| | | |
|---|---|---|
| C0040 | TetR | atgtccagattagataaaagtaaagtgattaacagcgcattag<br>agctgcttaatgaggtcggaatcgaaggtttaacaacccgtaa<br>actcgcccagaagctaggtgtagagcagcctacattgtattggc<br>atgtaaaaaataagcgggctttgctcgacgccttagccattgag<br>atgttagataggcaccatactcacttttgccctttagaaggggaa<br>agctggcaagattttttacgtaataacgctaaaagttttagatgtg<br>ctttactaagtcatcgcgatggagcaaaagtacatttaggtaca<br>cggcctacagaaaaacagtatgaaactctcgaaaatcaatta<br>gccttttatgccaacaaggtttttcactagagaatgcattatatgc<br>actcagcgctgtggggcattttactttaggttgcgtattggaagat<br>caagagcatcaagtcgctaaagaagaaagggaaacacctа<br>ctactgatagtatgccgccattattacgacaagctatcgaattatt<br>tgatcaccaaggtgcagagccagccttcttattcggccttgaatt<br>gatcatatgcggattagaaaaacaacttaaatgtgaaagtggg<br>tccgctgcaaacgacgaaaactacgctttagtagcttaataaca<br>ctgatagtgctagtgtagatcactaa |
| B0015 | Double terminator | ccaggcatcaaataaaacgaaaggctcagtcgaaagactgg<br>gcctttcgttttatctgttgtttgtcggtgaacgctctctactagagtc<br>acactggctcaccttcgggtgggcctttctgcgtttata |
| B0034 | Strong Ribosome Binding Site (RBS) | aaagaggagaaa |
| C0080<br>*(lva tag removed)* | AraC | atggctgaagcgcaaaatgatcccctgctgccgggatactcgtt<br>taacgcccatctggtggcgggtttaacgccgattgaggccaatg<br>gttatctcgatttttttatcgaccgaccgctgggaatgaaaggttat<br>attctcaatctcaccattcgcggtcaggggggtggtgaaaaatca<br>gggacgagaatttgtctgccgaccgggtgatattttgctgttccc<br>gccaggagagattcatcactacggtcgtcatccggaggctcgc<br>gaatggtatcaccagtgggtttactttcgtccgcgcgcctactgg<br>catgaatggcttaactggccgtcaatatttgccaatacgggtttct<br>ttcgcccggatgaagcgcaccagccgcatttcagcgacctgttt<br>gggcaaatcattaacgccgggcaaggggaagggcgctattc<br>ggagctgctggcgataaatctgcttgagcaattgttactgcggc<br>gcatggaagcgattaacgagtcgctccatccaccgatggataa<br>tcgggtacgcgaggcttgtcagtacatcagcgatcacctggca<br>gacagcaattttgatatcgccagcgtcgcacagcatgtttgcctg<br>tcgccgtcgcgtctgtcacatcttttccgccagcagttagggatta<br>gcgtcttaagctggcgcgaggaccaacgcatcagccaggcg<br>aagctgcttttgagcactacccggatgcctatcgccaccgtcgg<br>tcgcaatgttggttttgacgatcaactctatttctcgcgagtatttaa<br>aaaatgcaccggggccagcccgagcgagttccgtgccggttg<br>tgaagaaaaagtgaatgatgtagccgtcaagttgtcataa |
| C0061 | LuxI | atgactataatgataaaaaaatcggatttttggcaattccatcgg<br>aggagtataaaggtattctaagtcttcgttatcaagtgtttaagca<br>aagacttgagtgggacttagttgtagaaaataaccttgaatcag<br>atgagtatgataactcaaatgcagaatatatttatgcttgtgatga<br>tactgaaaatgtaagtggatgctggcgtttattacctacaacagg<br>tgattatatgctgaaaagtgtttttcctgaattgcttggtcaacaga<br>gtgctcccaaagatcctaatatagtcgaattaagtcgttttgctgt<br>aggtaaaaatagctcaaagataaataactctgctagtgaaatt<br>acaatgaaactatttgaagctatatataaacacgctgttagtcaa |

| | | |
|---|---|---|
| | | ggtattacagaatatgtaacagtaacatcaacagcaatagagc<br>gatttttaaagcgtattaaagttccttgtcatcgtattggagacaaa<br>gaaattcatgtattaggtgatactaaatcggttgtattgtctatgcct<br>attaatgaacagtttaaaaaagcagtcttaaatgctgcaaacga<br>cgaaaactacgctttagtagcttaa |
| C0062<br>*(S116A and M135I mutations were added)* | A variant of LuxR – LuxRG2C | atgaaaaacataaatgccgacgacacatacagaataattaat<br>aaaattaaagcttgtagaagcaataatgatattaatcaatgctta<br>tctgatatgactaaaatggtacattgtgaatattatttactcgcgat<br>catttatcctcattctatggttaaatctgatatttcaatcctagataat<br>taccctaaaaaatggaggcaatattatgatgacgctaatttaata<br>aaatatgatcctatagtagattattctaactccaatcattcaccaa<br>ttaattggaatatatttgaaaacaatgctgtaaataaaaaatctc<br>caaatgtaattaaagaagcgaaaacagcaggtcttatcactgg<br>gtttagtttccctattcatacggctaacaatggcttcggaatactta<br>gttttgcacattcagaaaaagacaactatatagatagtttattttta<br>catgcgtgtatgaacataccattaattgttccttctctagttgataat<br>tatcgaaaaataaatatagcaaataataaatcaaacaacgatt<br>taaccaaaagagaaaaagaatgtttagcgtgggcatgcgaa<br>ggaaaaagctcttgggatatttcaaaaatattaggttgcagtga<br>gcgtactgtcactttccatttaaccaatgcgcaaatgaaactcaa<br>tacaacaaaccgctgccaaagtatttctaaagcaattttaacag<br>gagcaattgattgcccatactttaaaaattaa |

**Supplemental Table 2. Gene circuits used in the study**

| Name | Gene construct |
|---|---|
| DSAC-T | K206000 + B0034 + GFP-lva + B0015 + K206000 + B0034 + araC + B0015 + K206000 + sg-pBAD + B0015 + Plux9 + B0034 + RFP-lva + B0015 + Plux9 + B0034 + luxRG2C + B0015 + Plux9 + sg-pLux9 + B0015 + J23117 + B0034 + TetR + B0015 |
| DSAC-C | K206000 + B0034 + GFP-lva + B0015 + K206000 + B0034 + araC + B0015 + K206000 + NT + B0015 + Plux9 + B0034 + RFP-lva + B0015 + Plux9 + B0034 + luxRG2C + B0015 + Plux9 + NT + B0015 + J23117 + B0034 + TetR + B0015 |
| CBS-T | K206000 + B0034 + GFP-lva + B0015 + K206000 + B0034 + araC + B0015 + K206000 + sg-pBAD + B0015 + Plux9 + B0034 + RFP-lva + B0015 + Plux9 + B0034 + luxRG2C + B0015 + Plux9 + sg-pLux9 + B0015 + J23117 + B0034 + TetR + B0015 + K206000 + B0034 + C0061 + B0015 + Plux9 + B0034 + araC + B0015 |
| CBS-C | K206000 + B0034 + GFP-lva + B0015 + K206000 + B0034 + araC + B0015 + K206000 + NT + B0015 + Plux9 + B0034 + RFP-lva + B0015 + Plux9 + B0034 + luxRG2C + B0015 + Plux9 + NT + B0015 + J23117 + B0034 + TetR + B0015 + K206000 + B0034 + C0061 + B0015 + Plux9 + B0034 + araC + B0015 |
| NCR cassette | P0440 + dCas9 + B0015 |

**Supplemental Table 3. Oligonucleotides used in this study**

| Sequence (5'→3') | Oligonucleotide description |
|---|---|
| ataagagttgcgataaaaagcgtg | Forward gRNA targeting pBAD |
| aaccacgctttttatcgcaactct | Reverse gRNA targeting pBAD |
| aaatgacacctgtaggatcgtac | Forward gRNA targeting pLux9 |
| aacgtacgatcctacaggtgtca | Reverse gRNA targeting pLux9 |
| tctggaattcgcggccgcttctagagtgacacctgtaggatcgtacaggg | Forward addition of TGAC to plux9 (EcoRI and XbaI) |
| ggactgcagcggccgctactagtacgctcttcttatggagaaacagtagagag | Reverse deletion of pBAD bases after TSS site (SapI, SpeI, and PstI) |
| ctagagtgacacctgtaggatcgtacagggttacgcaagaaaatggtttgttatagtcgaataaaagaagagcacaggctcttct | Forward Plux9 and non-targeting guide (SapI and SapI) |
| aacagaagagcctgtgctcttcttttattcgactataacaaaccattttcttgcgtaaccctgtacgatcctacaggtgtcact | Reverse Plux9 and non-targeting guide (SapI and SapI) |
| atgctgatttgtttttggcagc | Forward dCas9 specific primer |
| actccttggagaatccgcct | Reverse dCas9 specific primer |
| tctggaattcgcggccgcttctagagaaagaggagaaaggatctat | Forward addition of BioBrick restriction sites to dCas9 (EcoRI and XbaI) |
| ggactgcagcggccgctactagtacgcagaaaggcccacccgaa | Reverse addition of BioBrick restriction sites to dCas9 (SpeI and PstI) |
| ggactgcagcggccgctactagtattattaagctactaaagcgtagttttcgtcgtttgcagcagcaccggtggagtgacgac | Addition of lva tag to RFP (SpeI and PstI) |
| tctagaacaggctcttctgttttagagctagaaatagc | Forward dCas9 gRNA scaffold (XbaI and SapI) |
| ggactgcagcggccgctactagtagcaccgactcggtgccactt | Reverse addition of BioBrick restriction sites to dCas9 gRNA scaffold (SpeI and PstI) |
